# Modeling transcriptional profiles of gene perturbation with deep neural network

**DOI:** 10.1101/2021.07.15.452534

**Authors:** Wenke Liu, Xuya Wang, D R Mani, David Fenyö

## Abstract

**Background:** Cell line perturbation data could be utilized as a reference for inferring underlying molecular processes in new gene expression profiles. It is important to develop accurate and computationally efficient algorithms to exploit biological knowledge in the growing compendium of existing perturbation data and harness these for new predictions.

**Results:** We reframed the problem of inferring possible gene perturbation based on a reference perturbation database into a classification task and evaluated the application of deep neural network models to address this problem. Our results showed that a fully-connected multi-layer neural network was able to achieve up to 74.9% accuracy in a holdout test set, but the model generalizability was limited by consistency between training and testing data.

**Conclusion:** Capacity and flexibility enables neural network models to efficiently represent transcriptomic features associated with single gene knockdown perturbations. With consistent signals between training and testing sets, neural networks may be trained to classify new samples to experimentally confirmed molecular phenotypes.

## Background

As part of the NIH Library of Integrated Network-Based Cellular Signatures (LINCS) initiative, the Connectivity Map (CMap) dataset is a massive collection of transcriptomic profiles of genetically and pharmacologically perturbed human cell lines, obtained with the L1000 platform [1]. CMap provides a valuable reference for phenotypic manifestation of targeted perturbations, and the causal knowledge embedded in the compendium could aid clinical diagnosis as well as scientific discovery.

The CMap dataset is designed to serve as a lookup table, such that a query profile with unknown functional perturbation could be comprehensively compared to profiles in the dataset using a similarity measure. The CMap profiles thus identified as similar to the query, with known genetic and pharmacologic annotation, will provide insights into cellular and molecular mechanisms underlying the query profile. To overcome the off-target effect inherent to the short hairpin RNA (shRNA) knockdown technique, Consensus Gene Signatures (CGS) were calculated as weighted sum of raw profiles targeting the same gene, and it is advised to query the dataset on the CGS level with a nonparametric enrichment test [1].

From an algorithmic perspective, inferring the possible functional disruption in a transcriptomic profile by comparing it to perturbation signatures curated in the CMap dataset is similar to a prototype matching method. The problem can also be conceptualized as a classification task. In addition to direct query, one can also build a machine learning model that recognizes the transcriptome-wide effect of gene perturbations. In this light, the CMap profiles are training examples, with the known perturbation target genes corresponding to true labels. In addition to the established query method, application of state of the art machine learning algorithms to this task could improve the efficiency and accuracy of perturbation inference and facilitate downstream interpretation.

In the recent decade, deep learning methods have achieved stellar performance in a variety of tasks, ranging from image recognition to natural language processing [2]. This powerful technique also finds exciting applications in the biomedical field such as pathological image classification and transcription factor binding prediction [3, 4]. The method has also been used for gene expression inference. A three-layer neural network was utilized by the LINCS program to infer the full transcriptome from L1000 landmark genes and consistently exhibited better performance than linear regression, demonstrating the suitability of deep neural networks in modeling transcriptomic data [5].

The large model capacity of neural networks is especially suited for our classification task, where differences between gene perturbation groups lie in the high dimensional space of expression measurements and could be elusive to small scale, lower capacity models. When used to train a large scale neural network model the complex features of data can be encapsulated in the weights of the network and represented as nonlinear functions. At inference time, prediction for a given test sample would only require a single ‘forward propagation’ across the trained network, without direct comparison with the training examples. The multi-layer or ‘deep’ structure of neural network entails hierarchical abstraction of input measurements, and such abstractive, meta-gene level information may be utilized as shared inputs to prediction nodes for individual perturbation labels.

Here we report a neural network approach to the perturbation inference task described above. A deep neural network model was built using CMap profiles as training examples, and the model performance was benchmarked with both cross validation and external testing data. The aim of the current work is to assess that given a transcriptomic profile, whether DNN would facilitate the process of extracting relevant biological information from the CMap data. It is, to the best of our knowledge, the first attempt to explore an alternative to the CGS query method to identify gene knockdown using transcriptome profiles. Our results show that neural network models have great potential in predicting biological phenotypes from transcriptomic profiles, but the utility of such models rely on the consistency between training and testing data.

## Results

### Neural network model predicts gene perturbation with high accuracy in holdout test set

We first built a baseline model with only one output layer, which carried out softmax regression directly on the input data. This model is the generalization of logistic regression in the multi-class setting, and is implemented as a simple neural network with the input layer and the softmax output layer only. For each testing sample, the trained model inferred a vector of softmax probabilities for all 4314 possible gene categories. As the training and the testing sets are from the same population of transcriptomic profiles, concordance between model-inferred probabilities and the true label would indicate the extent to which the model captured complex patterns associated with different gene perturbation. When evaluated on holdout tests randomly sampled from the same full shRNA dataset as the training set, the baseline model showed adequate area under the receiver operating characteristic curve (AUROC) averaging on all classes, but the overall accuracy and average precision (AP) were relatively low (rank-1 accuracy 40.17±0.26%; rank-5 accuracy 57.93±0.05%; AP 0.4438±0.0011; average AUROC 0.9498±0.0005, mean ± standard error), suggesting that high model capacity is necessary.

Next, a deep neural network (DNN) model with stacked fully-connected layers was built following a heuristically optimized architecture (Fig. 1, Methods) with a bottleneck layer[6]. The same set of performance metrics were calculated for the DNN model (rank-1 accuracy 74.88±0.65%; rank-5 accuracy 83.41±0.48%; AP 0.7731±0.0062; average AUROC 0.9937±0.0001, mean ± standard error). The deep neural network showed significantly improved performance compared to the baseline model (*P* < 0.001, paired t-test on all metrics).

**Figure 1:**
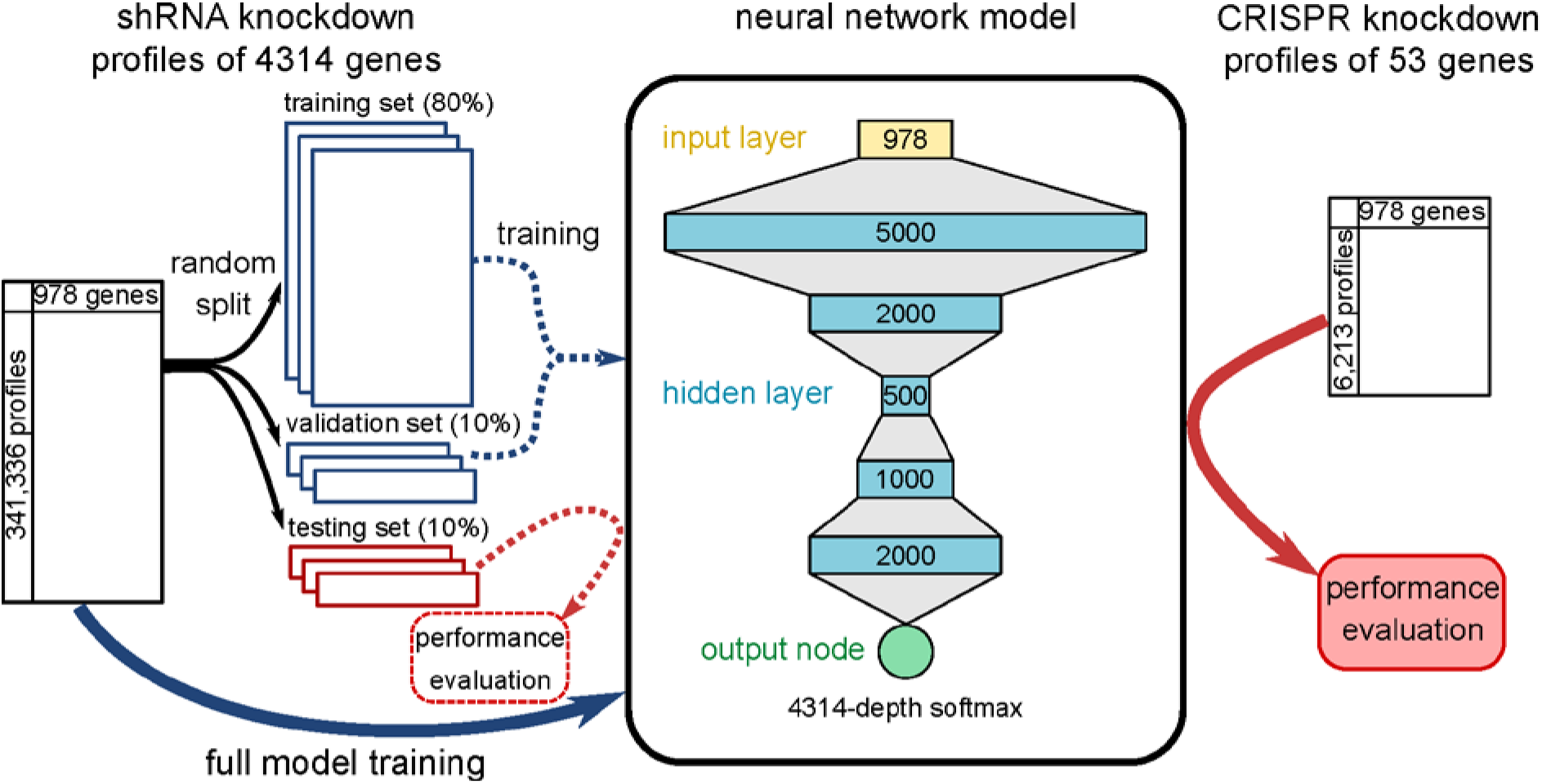
workflow of model training and testing. Neural network models was first trained on 80% of shRNA knockdown profiles and evaluated on holdout test sets. A final model was then trained with all shRNA data and tested on external test set of CRISPR knockdown profiles.

Prediction accuracy broken down by gene perturbations showed large variance. Genes with low accuracy had small training set size, suggesting that poor model performance for these genes resulted from underfitting due to lack of data (Fig. 2A). Out of the 4313 perturbed genes present in the test set, 946 are landmark genes, and 3367 are non-landmark genes. Group averaged accuracy of the landmark genes was higher than the non-landmark category (two-tail unpaired t-test, rank-1, *P*=0.0041; rank-5, *P*=0.0002), possibly resulting from the fact that landmark genes were directly measured, and hence functioned as more robust input feature values (Fig. 2B).

**Figure 2:**
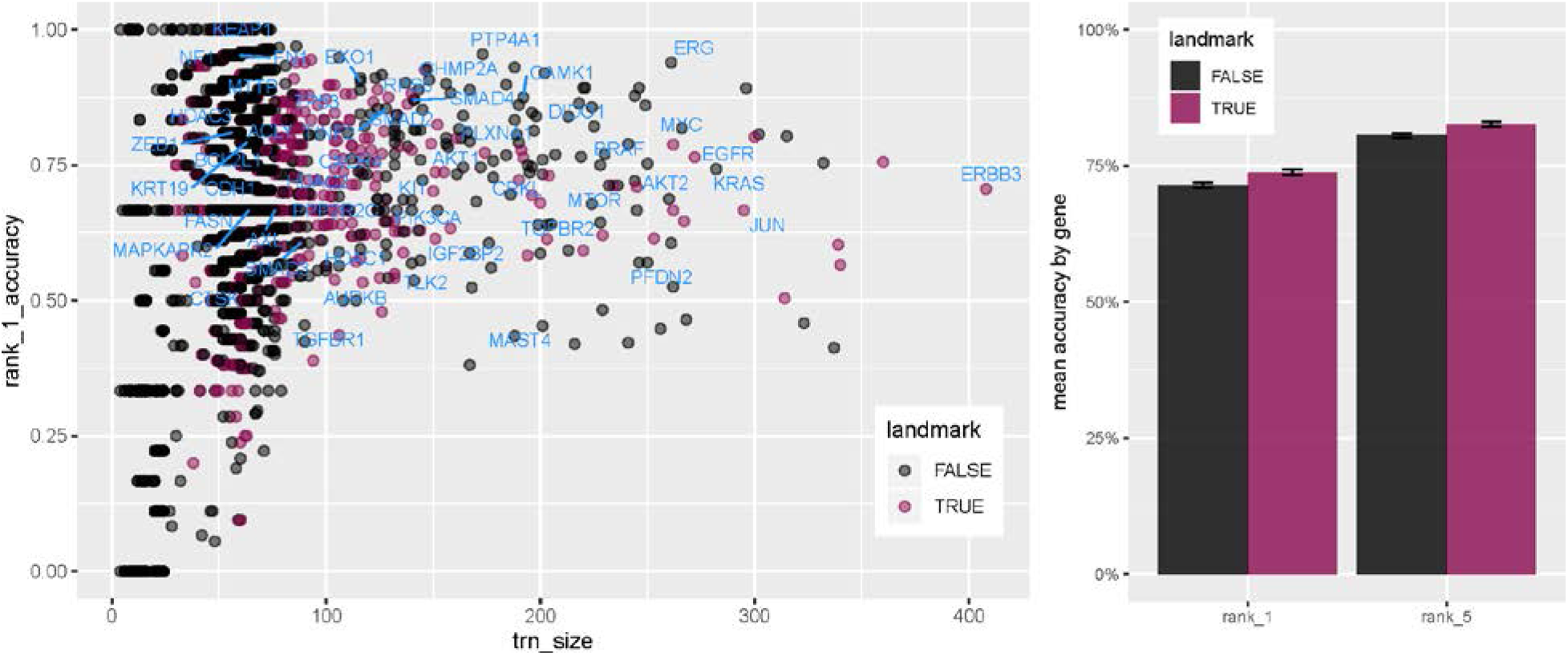
Prediction accuracy of neural network model on holdout test set. A, across-split average rank-1 accuracy for 4314 genes plotted against number of training examples. Genes for which knockdown data was also available in the CRISPR data set are labeled in blue text. B, group average rank-1 and rank-5 accuracy values stratified by whether the perturbed genes were directly measured as a landmark gene.

### Neural network model does not generalize across platforms

To further assess generalizability of the trained model, we took advantage of the Clustered Regularly Interspaced Short Palindromic Repeats (CRISPR) knockdown subset of CMap data characterized on the same L1000 platform. 50 of the 53 CRISPR knockdown genes were also perturbed by shRNA. 5884 profiles with true labels among the 50 genes were scored using the final shRNA-trained model using all 341,336 profiles grouped into 4314 gene knockdown categories. The overall rank-1accuracy (4.47%), rank-5 accuracy (6.70%), AP (0.1031) and average AUROC (0.6078) evaluated on the CRISPR test set were significantly lower than that obtained on the holdout set reported above, but still higher than that of the baseline softmax regression model (rank-1 accuracy 2.79%; rank-5 accuracy 5.49%; AP 0.0787, average AUROC 0.5901). Only 6 out of the 50 genes (*KEAP1*, *RPS6*, *DIDO1*, *MYC*, *MTOR*, *BRAF*) exhibited rank-1 accuracy higher than 10% (Fig. 3A).

**Figure 3:**
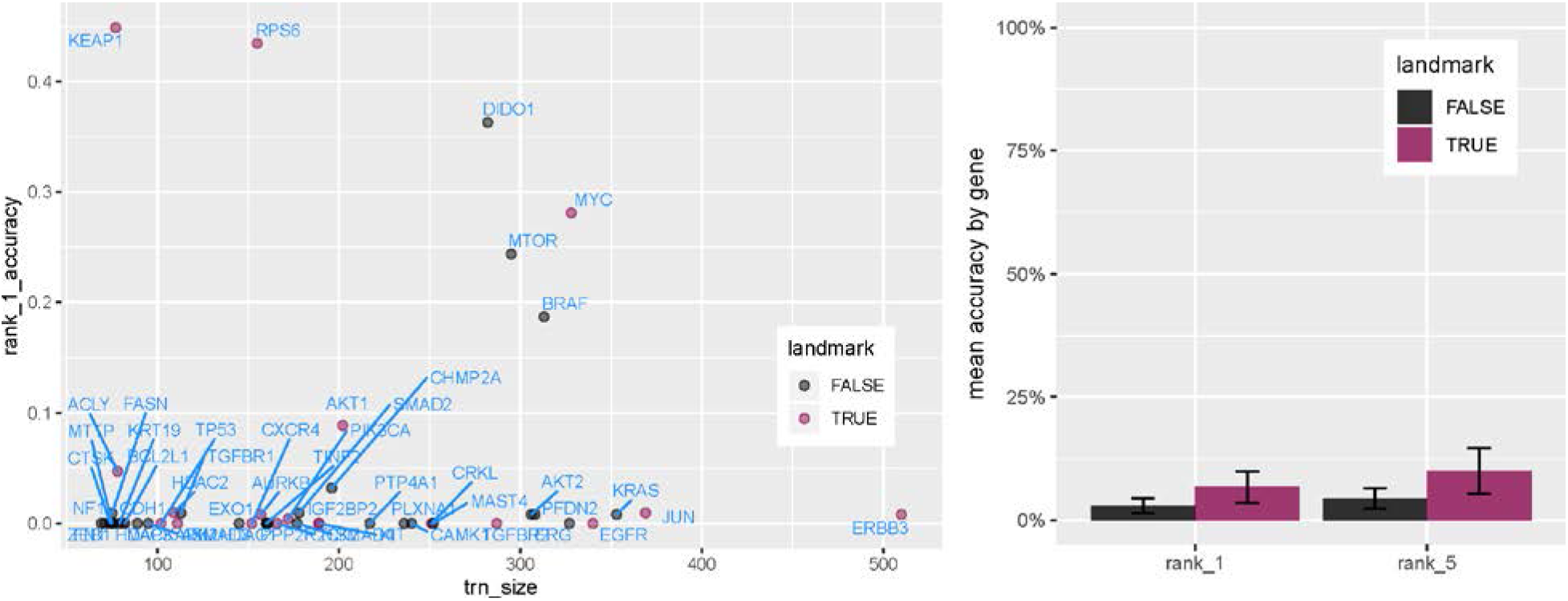
Prediction accuracy of neural network model on external test set. A, rank-1 accuracy for 50 genes shared between shRNA and CRISPR test set plotted against number of examples used to train the final model. B, group average rank-1 and rank-5 accuracy values stratified by whether the perturbed genes were directly measured as a landmark gene.

Consistent with the shRNA holdout test (Fig. 3B), the average accuracy of landmark genes (n=20; rank-1, 6.7±3.2%; rank-5, 10.0 ±4.6%) in the CRISPR dataset was higher than that of non-landmark genes (n=30; rank-1, 2.9±1.5%; rank-5, 4.4 ±2.1%), but the differences were not statistically significant (two-tail unpaired t-test, rank-1, *P*=0.25; rank-5, *P*=0.22).

Given the poor performance of the shRNA data trained model on CRISPR data, both model adequacy and data distribution consistency were at question. We further investigated this issue by comparing the neural network model to the established query method. In addition to transcriptomic profiles of experimental samples, CMap shRNA data also contained consensus gene signatures (CGS) of 33,839 combinations of cell lines and perturbed genes. The CGS was calculated as weighted averages of profiles with the same cell line and gene target condition, providing a representation of the gene expression information associated with the specific context and treatment. Systematic comparison between a new input profile and all CGSs therefore serve as a prototype classifier, by which the most probable gene perturbation in the query sample can be inferred. An enrichment score based similarity measure has been recommended to map the ‘connectivity’ query and expression signatures [1].

We constructed a pair of up-down gene sets from each of the 5884 CRISPR profiles and calculated a vector of weighted connectivity scores (WTCS), which indicated similarity of the query profile to each of the 33,839 CGSs. After aggregation of scores associated with the same gene, the 4318 element vector can be rearranged into a ranked list of all targeted genes, which can be further transformed into softmax probabilities comparable to that produced by the DNN model. The prediction performance of CGS query method across 5884 profiles was also unsatisfactory (rank-1 accuracy 2.75%; rank-5 accuracy 4.95%; AP 0.0975, average AUROC 0.5927), flagging the same model generalization problem, indicating that shRNA CGSs shared only limited features with CRISPR profiles under the same gene perturbation category.

Number of samples in which the true gene target was ranked at the top was 162 for the CGS query and 263 for the neural network model, with 98 correctly predicted perturbations shared by both methods (Fig. 4B). Breakdown of true-positive predictions by targeted genes further showed that the two methods may exploit different information, as the true positive cases did not overlap within the same gene (Fig. 4C).

**Figure 4:**
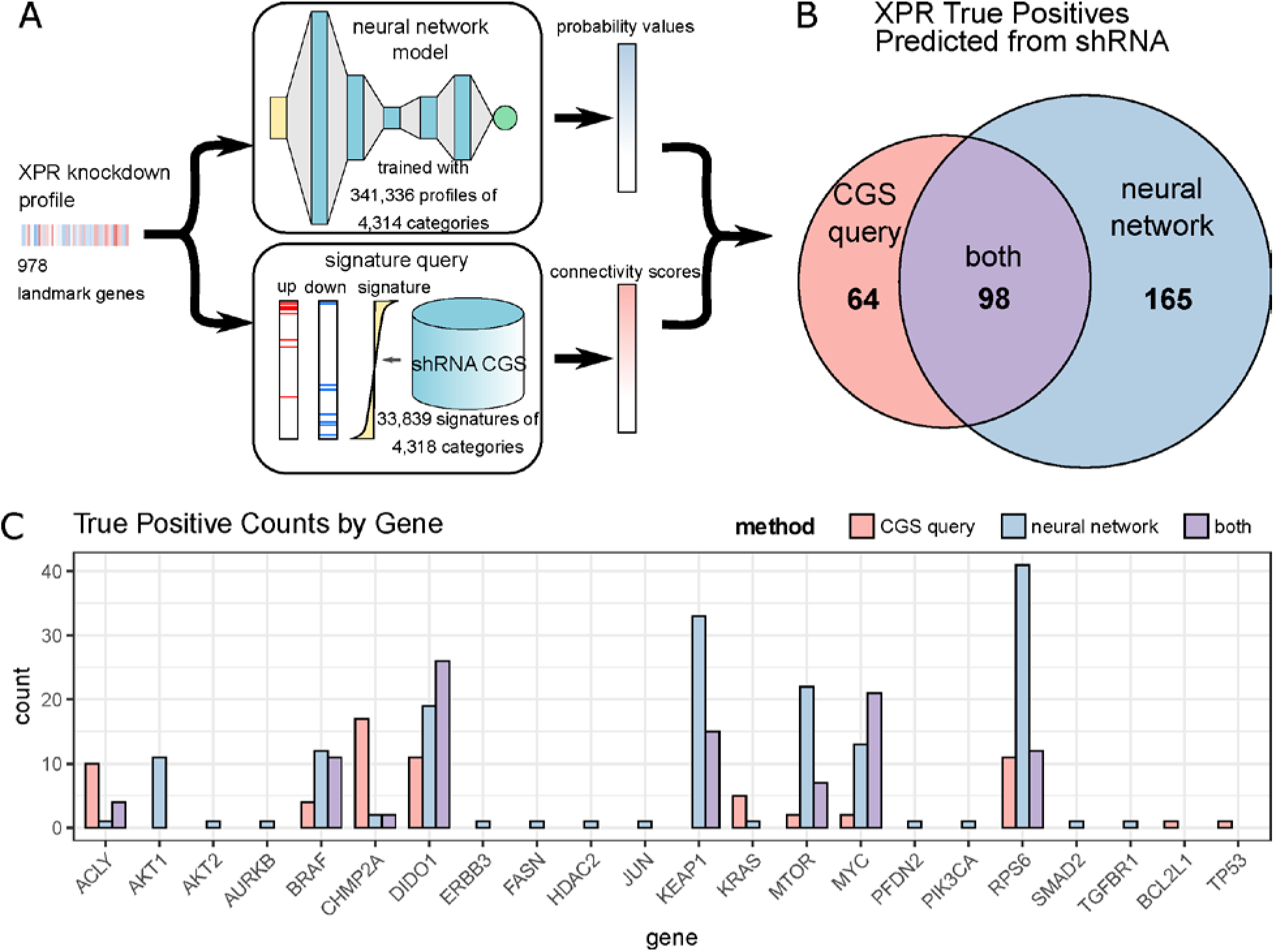
Performance comparison between neural network method and CGS query method. A, schematic of the workflow comparing two methods. Test CRISPR knockdown profiles were submitted to both the neural network model and consensus gene signature query, each giving rise to a ranked list of scores corresponding to gene labels. B, Venn diagram of test profiles for which true labels were assigned the largest score by the two methods. C, Gene-level breakdown of test sample counts that are correctly predicted by CGS only (red), neural network only (blue) or both methods (purple).

## Discussion

Our results showed that a multi-layer neural network model can recognize the complex transcriptomic patterns associated with specific gene knockdowns in a defined technical context. When the training and the testing data were sampled from the same population, the model could achieve 74.9% accuracy in a multi-class classification task with 4314 labels. However, the model trained with shRNA knockdown profiles was not generalizable to data generated by a CRISPR knockdown system. Consensus gene signature (CGS) query method applied to the same testing data also gave rise to similarly unsatisfactory accuracy, suggesting that data inconsistency is the primary cause for lack of model generalizability.

The consistency between the neural network model and the CGS query predictions suggested that noise due to different knockdown systems overwhelmed common biological signal. In fact, it has been independently reported that cross query similarity between shRNA and CRISPR CGS showed only modest connectivity [7]. Therefore, the poor generalizability of neural network predictions is more likely due to systematic differences between the shRNA and CRISPR datasets, rather than model capacity.

## Conclusions

Neural networks are powerful models that have been successfully applied to biomedical data. We showed that a deep neural network with stacked fully-connected layers was able to capture complex patterns in transcriptome profiles generated by perturbation using the same experimental methodology. However, the ability of such nonlinear hierarchical models to capture true biological information is severely limited by the wide variation in molecular change induced by changes in the experimental procedure used for perturbation. Neural networks can relate complex biological information in new testing data to common features found in training data, when no systematic difference is present between the two data sets.

## Methods

### Data

In this study we utilized the current LINCS data release (GEO accession number: GSE106127) [1, 7]. The shRNA genetic knockdown subset of level-4 CMap data was comprised of 334,648 profiles for 4,314 knockdown genes. Each level-4 profile contains differential expression values of 978 landmark genes measured in one sample, calculated as z-scores with respect to the entire population of the experimental plate. Profiles with the same knockdown label perturb the same gene, but the samples were obtained from different cell lines, treated with different hairpin sequences and measured within different replicates. The CRISPR knockdown data was profiled on the same platform, containing 6,213 level-4 profiles for 53 knockdown genes, most of which are also perturbed with the shRNA method (5884 profiles of 50 genes). The 33,839 consensus gene signatures (CGS) were collapsed from level-5 shRNA data, with each signature representing a gene perturbation in a specific cell line. Vector controls were excluded from all datasets. For model training, the tuple of landmark gene measurements and perturbation label for all samples were aggregated as a dataset, randomly shuffled and batched.

### Neural network model

Neural network architecture design was performed under Bayesian optimization framework with the GPyOpt software [8]. Search over a discrete hyper-parameter space of layer sizes (5000, 2000, 1000, 500, 200, 100), learning rate (10^−4^, 10^−5^) and lambda of L_2_ regularization (10^−3^, 10^−4^) with accuracy on validation set as the target did not reach consistent local optima on different random data splits. The final model was built in a heuristic manner with the following architecture: input layer with 978 nodes, each corresponding to one landmark gene; 5 hidden layers with 5000, 2000,500, 1000, 2000 nodes; final node outputs a softmax vector of depth 4314 representing probability values of an input profile belonging to each of the 4314 gene perturbation categories. All hidden layers are fully-connected with exponential linear unit (ELU) as node activation function, and cross-entropy between predicted probability and one-hot encoded true label was used as the loss function. Models were trained for 100 epochs with Adam optimizer and batch size of 128, with dropout probability of 0.1, at learning rate of 10^−4^ and L_2_ regularization lambda of 10^−4^. An ensemble of 5 models with the same hyper-parameters and different initiations were trained, and the output probabilities of 5 models were averaged at prediction time. The model was implemented with TensorFlow 1.12.0 and trained on Nvidia Quadro P600 GPU.

### Consensus gene signature (CGS) query

Query on the CGS data was based on enrichment score calculation described in Subramanian et al. In brief, for each query input (CRISPR knockdown profile), 50 genes with the highest and lowest expression among the 978 measured landmark genes were collected into two gene sets, q_up_ and q_down_. For each shRNA knockdown consensus signature r, enrichment scores (ES) of q_up_ and q_down_ were calculated as the maximum deviation from zero of the curve:

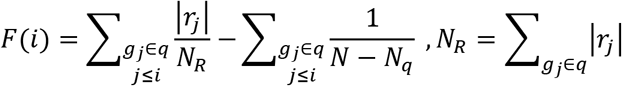

N and N_q_ are total number of genes in the signature and the query gene set, respectively. In this case, N=978 and N_q_=50. Enrichment scores were calculated using R package fGSEA [9]. Bi-directional weighted connectivity score (WTCS) was further calculated as:

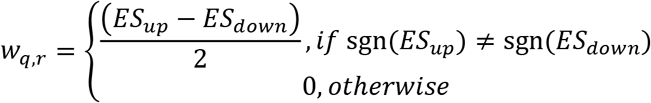

WTCS of each query compared to all CGSs are calculated, and scores obtained from same-gene perturbation in different cell line were aggregated by taking the maximum value.

### Performance evaluation

We assess the performance of the baseline model and the final neural network model on holdout testing sets. 334,648, shRNA knockdown profiles (excluding controls) were randomly split into 3 training (80%), validation (10%) and testing (10%) sets, stratified with respect to gene knockdown labels. On each data split, neural network model was trained on the training set, and model performance was monitored on-line with the validation set. At inference time, landmark gene expression values of test profiles were propagated into the trained model, and the predicted softmax probability vector for each gene category was obtained. Four metrics were calculated based on the predicted probabilities: rank-1 accuracy (fraction of test samples on which the model predicted the largest probability for the true class), rank-5 accuracy (true class among the top five predicted probabilities), average precision score (AP) with macro average, area under the receiver operating characteristic curve (AUROC) with macro average. For AP and AUROC, the ‘macro’ method calculates the metric for each class in a one-vs-other manner and takes the average, regardless of the size of each class. This is in contrast to the accuracy metric that assigns higher weights to larger classes [10]. Mean metric values were aggregated across different random splits.

To test the generalizability of the neural network model, a full model of the same architecture was trained with all shRNA profiles and evaluated on 5884 CRISPR knockdown profiles that share perturbation targets with the shRNA dataset. General workflow is illustrated in Fig. 1.Gene-level predictions were made for the same 5884 CRISPR profiles using the CGS query method as described above. Enrichment scores for 4212 categories common between the two methods were transformed with the softmax function, resulting in a probability vector for each test sample, in the same format as the DNN predictions. The same set of four metrics (rank-1 and rank-5 accuracy, AP, AUROC) as well as true positive counts were calculated for both methods.

## List of Abbreviations

CMap: Connectivity Map
DNN: Deep Neural Network
shRNA: short hairpin RNA
CRISPR: Clustered Regularly Interspaced Short Palindromic Repeats
CGS: Consensus Gene Signatures
ES: Enrichment Score
WTCS: Weighted Connectivity Score
ELU: Exponential Linear Unit

## Declarations

### Ethics approval and consent to participate

Not applicable

### Consent for publication

Not applicable

### Availability of data and materials

The data analyzed in the current study was previously published [7], available in the GEO repository: https://www.ncbi.nlm.nih.gov/geo/query/acc.cgi?acc=GSE106127

### Authors’ contributions

DF and DRM designed the project. WL and XW performed the analysis. WL wrote the paper. All authors discussed the results and approved the final draft.

## Acknowledgements

Not applicable

## Authors’ information (optional)

Not applicable

